# Does the human placenta express the canonical cell entry mediators for SARS-CoV-2?

**DOI:** 10.1101/2020.05.18.101485

**Authors:** Roger Pique-Regi, Roberto Romero, Adi L. Tarca, Francesca Luca, Yi Xu, Adnan Alazizi, Yaozhu Leng, Chaur-Dong Hsu, Nardhy Gomez-Lopez

## Abstract

The pandemic of coronavirus disease 2019 (COVID-19) caused by the severe acute respiratory syndrome coronavirus 2 (SARS-CoV-2) has affected over 3.8 million people, including pregnant women. To date, no consistent evidence of vertical transmission for SARS-CoV-2 exists. This new coronavirus canonically utilizes the angiotensin-converting enzyme 2 (ACE2) receptor and the serine protease TMPRSS2 for cell entry. Herein, building upon our previous single cell study of the placenta (Pique-Regi, 2019), another study, and new single-cell/nuclei RNA-sequencing data, we investigated the expression of ACE2 and TMPRSS2 throughout pregnancy as well as in third-trimester chorioamniotic membranes. We report that co-transcription of ACE2 and TMPRSS2 is negligible, thus not a likely path of vertical transmission for SARS-CoV-2 at any stage of pregnancy. In contrast, receptors for Zika virus and cytomegalovirus which cause congenital infections are highly expressed by placental cell types. These data suggest that SARS-CoV-2 is unlikely to infect the human placenta through the canonical cell entry mediators; yet, other interacting proteins could still play a role in the viral infection.

## MAIN TEXT

The placenta serves as the lungs, gut, kidneys, and liver of the fetus [1, 2]. This fetal organ also has major endocrine actions that modulate maternal physiology [1] and, importantly, together with the extraplacental chorioamniotic membranes shield the fetus against microbes from hematogenous dissemination and from invading the amniotic cavity [3]. Indeed, most pathogens that cause hematogenous infections in the mother are not able to reach the fetus, which is largely due to the potent protective mechanisms provided by placental cells (i.e. trophoblast cells: syncytiotrophoblasts and cytotrophoblasts) [4]. Yet, some of these pathogens such as *Toxoplasma gondii*, Rubella virus, herpesvirus (HSV), cytomegalovirus (CMV), and Zika virus (ZIKV), among others, are capable of crossing the placenta and infecting the fetus, causing congenital disease [5, 6].

In December 2019, a local outbreak of pneumonia caused by a novel coronavirus—severe acute respiratory syndrome coronavirus 2 (SARS-CoV-2)—was reported in Wuhan (Hubei, China) [7]. After exposure to SARS-COV-2, susceptible individuals can develop coronavirus disease 2019 (COVID-19) consisting of symptoms that may range from fever and cough to severe respiratory illness; in some cases, COVID-19 is life-threatening [8, 9]. Since the onset of the outbreak, more than 3.8 million COVID-19 cases have been confirmed, accounting for more than 269,000 deaths [10]. This pandemic has now spread throughout the entire world with recent epicenters in Europe (Italy and Spain) and the United States. By April 2019, the states of New York and Michigan were the most affected [10], given that the metropolitan areas of New York City and Detroit possess a large population subject to health disparities such as limited access to health care, chronic exposure to pollution, and pre-existing cardiovascular conditions [11].

Pregnant women and their fetuses represent a high-risk population in light of the COVID-19 outbreak [12-20] since viral infections such as influenza [21-27], varicella [28-32], Ebola [33, 34], and measles [35, 36] show increased severity in this physiological state. Other coronavirus such as SARS-CoV-1 and MERS-CoV have severe effects to both the mother and the fetus, but vertical transmission has not been proven [37-41] albeit these studies included very few number of cases. In contrast with the abovementioned viral infections, only ∼15% of pregnant women test positive for SARS-CoV-2 and a small fraction of them are symptomatic [42], most of whom (92%) experience only a mild illness [43]. Thus, the clinical characteristics of pregnant women with COVID-19 appear similar to those of non-pregnant adults [44]. In addition, thus far, no conclusive evidence of vertical transmission has been generated [45-49]. Consistently, infants born to mothers with COVID-19 test negative for SARS-CoV-2, do not develop serious clinical symptoms (e.g., fever, cough, diarrhea, or abnormal radiologic or hematologic evidence), and are promptly discharged from the hospital [50]. Nevertheless, new evidence has emerged suggesting that the fetus can respond to SARS-CoV-2 infection.

Case reports showed that a small fraction of neonates born to women with COVID-19 tested positive for the virus at 1-4 days of life [51, 52]; yet, these neonates subsequently tested negative on day 6-7 [51]. In addition, serological studies revealed that a few neonates born to mothers with COVID-19 had increased concentrations of SARS-CoV-2 immunoglobulin (Ig)M as well as IgG [53, 54]. The elevated concentrations of IgG are likely due to the passive transfer of this immunoglobulin from the mother to the fetus across the placenta. However, the increased levels of IgM suggest that the fetus was infected with SARS-CoV-2 since this immunoglobulin cannot cross the placenta due to its large molecular weight. Nonetheless, all neonates included in the above-mentioned studies tested negative for the virus and did not present any symptoms [53, 54].

More recently, two case reports have been reported where SARS-CoV-2 RNA has been detected in amniotic fluid and placental tissues. In the first case report, the viral RNA was detected in amniotic fluid from a woman who was severely affected and died of COVID-19 [55]. The premature neonate tested negative for SARS-CoV-2 after delivery but 24hr later tested positive [55]. In the second case report, the viral RNA was detected in the placenta and umbilical cord from a woman with severe pre-eclampsia, placental abruption, and other complications; yet, none of the fetal tissues tested positive [56]. Therefore, whether SARS-CoV-2 can reach the fetus by crossing the placenta is still unclear.

Cell entry and spreading of SARS-CoV-2 has been widely thought to depend on the angiotensin-converting enzyme 2 (ACE2) receptor [57, 58] and the serine protease TMPRSS2 [59]. In the study herein, we investigated whether the receptors responsible for SARS-CoV-2 infection are expressed in the human placenta (including the decidual tissues) throughout the three trimesters of pregnancy using publicly available single-cell RNA-sequencing (scRNA-seq) data [60, 61] together with newly generated data (Table S1).

Strikingly, we found that very few cells co-express ACE2 and TMPRSS2 **(Fig. 1A and B)**. Using a very permissive threshold of expression of one transcript per cell, only four cells with co-expression were detected in any of the three trimesters, resulting in an estimated < 1/10,000 cells. Our first-trimester data are in agreement with a prior report showing that there is minimal expression of ACE2 at the human maternal-fetal interface [62]; however, the same dataset was recently used to report the opposite [63]. Nonetheless, the co-expression of ACE2 and TMPRSS2 was not examined by either study. We also evaluated the expression of SARS-CoV2 receptors in the chorioamniotic membranes (also known as the extraplacental membranes) in the third trimester, since these tissues may also serve as a point of entry for microbial invasion of the amniotic cavity and potentially the fetus [64]. Again, co-expression of ACE2 and TMPRSS2 was minimally detected in the chorioamniotic membranes **(Fig. 1A and B)**.

**Figure 1.**
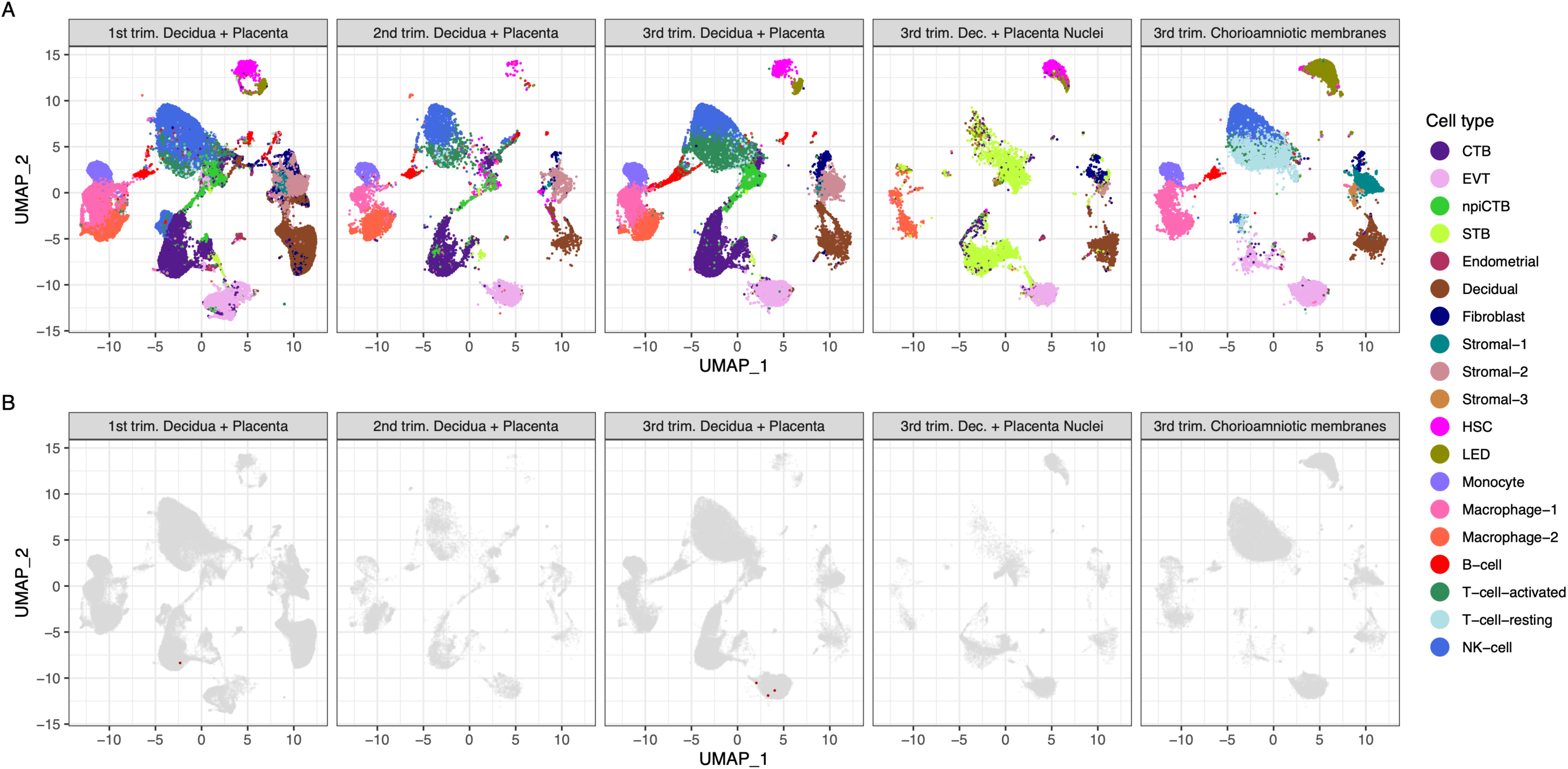
Transcriptional map of the human placenta, including the decidua, in the three trimesters of pregnancy. **A**. Uniform Manifold Approximation Plot (UMAP), where dots represent single cells/nuclei and are colored by cell type (abbreviations used are: STB, Syncytiotrophoblast; EVT, Extravillous trophoblast; CTB, cytotrophoblast; HSC, hematopoietic stem cell; npiCTB, non proliferative interstitial cytotrophoblast; LED, lymphoid endothelial decidual cell) **B**. UMAP plot with cells/nuclei co-expressing one or more transcripts for ACE2 and TMPRSS2, genes that are necessary for SARS-CoV-2 viral infection and spreading, in red.

A challenge in scRNA-seq studies is generating high-quality single-cell suspensions containing both rare and difficult-to-dissociate (e.g. multinucleated cells) cell types. This is likely the reason why the reported scRNA-seq studies of the human placenta contain a low fraction of syncytiotrophoblast cells [STB, multinucleated cells forming the outermost fetal component of the placenta in direct contact with the maternal circulation (i.e., intervillous space)] [60, 61, 65]. Therefore, we considered whether the expression of ACE2 and TMPRSS2 was minimally observed in the placental cell types due to the reduced fraction of STB cells (i.e., dissociation bias). To address this possibility, we prepared single-nucleus suspensions of the placental tissues (including the decidua basalis) and performed single-nuclear RNAseq (snRNA-seq), which reduces the dissociation bias against large cells [66]. An important advantage of snRNA-seq is its compatibility with biobank frozen samples; therefore, we pooled 32 placental villi/decidua samples collected in the third trimester (Table S2). This represents the first snRNA-seq study of the placental tissues. A limitation of snRNA-seq is that it has a higher background compared to scRNAseq but this should not affect the analyses reported here. As expected, a larger fraction of STB cells/nuclei was observed using snRNA-seq compared to scRNA-seq **(Fig. 1A)**. Consistent with the scRNAseq analyses, the snRNAseq data demonstrated that co-expression of ACE2 and TMPRSS2 is unlikely in the placental tissues **(Fig. 1B)**. Finally, we explored the co-expression of ACE2 and TMPRSS2 in third-trimester placental tissues by mining two microarray datasets that we have previously reported [67, 68]. These analyses of bulk gene expression data revealed that while ACE2 was detected above background in most of the samples, TMPRSS2 was largely undetected **(Table S3)**. Collectively, these results consistently indicate that the human placental tissues negligibly co-express ACE2 and TMPRSS2. This reduced expression contrasts with the high expression of ACE2 and TMPRSS2 in nasal goblet and ciliated cells within the human airways, lungs, and gastrointestinal tract, which are targeted during COVID-19 [69-71]. Therefore, our results suggest that vertical transmission of SARS-CoV-2 is unlikely to occur unless facilitated by other concomitant pathological conditions resulting in a breach of the maternal-fetal crosstalk.

There is a possibility, however, that SARS-CoV-2 could infect the human placenta by using alternate entry routes by interacting with other proteins [72]. The expression of additional SARS-CoV-2-related receptors or proteins in the human placenta is shown in **Fig. 2, CoV-Alt**; however, further research is required to test their participation in the pathogenesis of COVID-19. For example, *in vitro* studies suggest that BSG (Basigin, also called CD147 or EMMPRIN, transmembrane glycoprotein belonging to the immunoglobulin superfamily) provides an alternate entry for SARS-CoV-2 when ACE2 and TMPRSS2 are not expressed [73-75]. We found that the placenta and chorioamniotic membranes expressed high levels of BSG throughout pregnancy **(Fig. 2, CoV-Alt)**; yet, this transcript is also widely expressed in all human tissues and cell types **(Fig. S1)**. Therefore, it is unlikely that this protein alone is a sufficient requirement for SARS-CoV-2 viral entry and other proteins may be required to explain the cell-type primarily affected by COVID-19. Moreover, cathepsin L (CSTL) and FURIN may also function as proteases priming the SARS-CoV-2 S protein [76]. We found that these proteases are highly expressed by the placental tissues throughout gestation **(Fig. 2, CoV-Alt).** Nevertheless, these proteases may not provide sufficient levels of priming by themselves [77-79] when tested with SARS-CoV-1, yet this has not been verified for SARS-CoV-2. Given that the placental tissues are enriched in maternal and fetal macrophages [61], and that a subset of these immune cells expressing sialoadhesin (SIGLEC1, also known as CD169) can contribute to viral spread during SARS-CoV-2 infection [80, 81], we also investigated the expression of SIGLEC1 in this study. As expected, SIGLEC1 was expressed by macrophages in the placenta and chorioamniotic membranes and, to a lesser extent, in T cells **(Fig. 2, CoV-Alt)**. However, even if the virus could infect the placental/decidual macrophages expressing SIGLEC1, this is not sufficient for viral spreading. The expression of ADAM17 was also investigated in the placental tissues since this metalloproteinase competes with TMPRSS2 in ACE2 processing [82]. The placenta and chorioamniotic membranes highly expressed ADAM17 **(Fig. S2)**; however, only cleavage by TMPRSS2 results in augmented SARS-S-driven cell entry [82]. While these CoV-Alt molecules may be used for SARS-CoV-2 infection, they are likely to be less efficient than ACE2 and TMPRSS2, which are already targeted for antiviral interventions [59]; yet, new candidate host:viral interacting proteins and possible drugs are being investigated [72].

**Figure 2.**
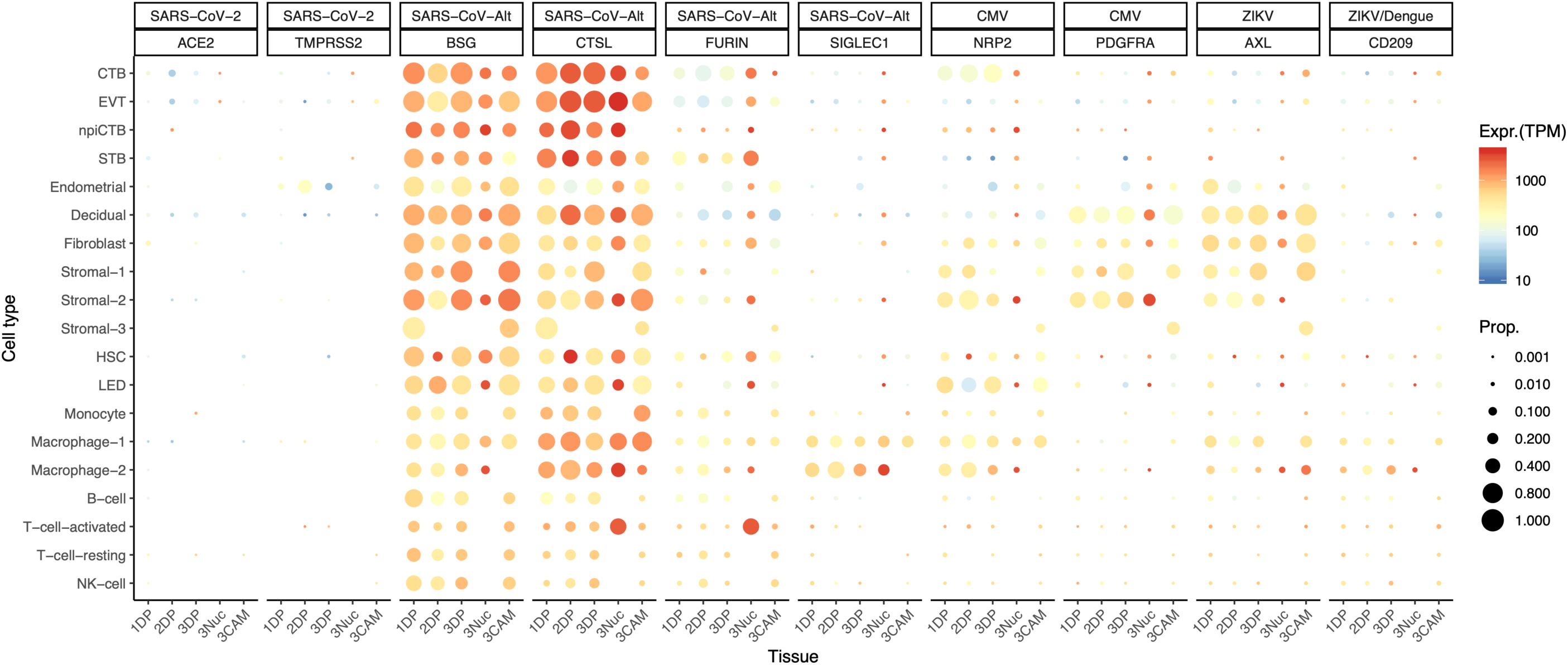
Dot plot depicting the expression of different viral receptors/molecules used by SARS-CoV-2, ZIKV, and CMV. Each row represents a different cell type, and columns are grouped first by virus type, receptor/molecule gene, and placental tissue/time-of sampling (1DP, 2DP and 3DP represent the first, second, and third trimester, 3Nuc represents the third trimester nuclei, and 3CAM represents the third trimester chorioamniotic membranes). The size of the dot represents the proportion of cells that express the receptor with more than zero transcripts, and the color represents the average gene expression for the subset of cells expressing that gene in transcripts per million (TPM). Cell type abbreviations used are: STB, Syncytiotrophoblast; EVT, Extravillous trophoblast; CTB, cytotrophoblast; HSC, hematopoietic stem cell; npiCTB, non proliferative interstitial cytotrophoblast; LED, lymphoid endothelial decidual cell.

Given that the main mediators for cell entry of SARS-CoV-2 were minimally expressed by the human placenta, we also investigated whether the receptors for congenital viruses such as CMV and ZIKV, which are known to infect and cross the placenta [5, 6], were detectable using our pipeline. Known receptors for CMV include NRP2 [83], PDFGRA [83], and CD46 [84]. Notably, all of these receptors were highly expressed in several placental cell types **(Fig. 2, CVM and Fig. S2)**. Next, we investigated the expression of the AXL receptor for ZIKV [85, 86] as well as other related molecules such as CD209 [87] and TYRO3 [88]. Consistent with vertical transmission, AXL, the preferred receptor for ZIKV, was highly expressed by the cells of the human placenta and chorioamniotic membranes throughout gestation **(Fig. 2, ZIKV).** The expression of CD206 was mainly found in the maternal and fetal macrophage subsets, as expected [89, 90]. Yet, the expression of TYRO3 was low **(Fig. S2)**, consistent with the view that TAM receptors are not essential for ZIKV infection [91]. The expression of other viral receptors involved in congenital disease was also documented in the placental tissues **(Fig. S2)**.

In conclusion, the single-cell transcriptomic analysis presented herein provides evidence that SARS-CoV-2 is unlikely to infect the placenta and fetus since its canonical receptor and protease, ACE2 and TMPRSS2, are only minimally expressed by the human placenta throughout pregnancy. In addition, we showed that the SARS-CoV-2 receptors are not expressed by the chorioamniotic membranes in the third trimester. However, viral receptors utilized by CMV, ZIKV, and others are highly expressed by the human placental tissues. While transcript levels do not always correlate with protein expression, our data indicates a low likelihood of placental infection and vertical transmission of SARS-CoV-2. However, it is still possible that the expression of these proteins is much higher in individuals with pregnancy complications related with the renin-angiotensin-aldosterone system, which can alter the expression of ACE2 [92]. The cellular receptors and mechanisms that could be exploited by SARS-CoV-2 are still under investigation [72]; yet, single-cell atlases can help to identify cell types with a similar transcriptional profile to those that are known to participate in COVID-19.

## METHODS

### Data availability

Placenta and decidua scRNA-seq data from first-trimester samples were downloaded through ArrayExpress (E-MTAB-6701). Data for third-trimester samples previously collected by our group are available through NIH dbGAP (accession number phs001886.v1.p1), and newly generated second-trimester scRNA-seq and third-trimester snRNA-seq data are being deposited in the same repository (Table S1). All software and R packages used herein are detailed in the “scRNA-seq and snRNA-seq data analysis.” Scripts detailing the analyses are also available at https://github.com/piquelab/sclabor.

### Sample collection and processing, single-cell/nuclei preparation, library preparation, and sequencing

#### Human subjects

Placental tissues were obtained immediately after a clinically indicated delivery from (i) a patient diagnosed with placenta accreta at 18 weeks of gestation and (ii) 32 patients spanning different conditions at the third trimester (Table S2). A sample of the basal plate of the placenta including the decidua basalis and placental villi tissue was (i) dissociated as previously described [61] for scRNA-seq or (ii) preserved in RNAlater and subsequently frozen for snRNA-seq. The collection and use of human materials for research purposes were approved by the Institutional Review Board of the Wayne State University School of Medicine. All participating women provided written informed consent prior to sample collection.

#### Single-cell preparation

Cells from the placental villi and basal plate were isolated by enzymatic digestion using previously described protocols with modifications [61, 65, 93]. Briefly, placental tissues were homogenized using a gentleMACS Dissociator (Miltenyi Biotec, San Diego, CA) either in an enzyme cocktail from the Umbilical Cord Dissociation Kit (Miltenyi Biotec) or in collagenase A (Sigma Aldrich, St. Louis, MO). After digestion, homogenized tissues were washed with ice-cold 1X phosphate-buffered saline (PBS) and filtered through a cell strainer (Fisher Scientific, Durham, NC). Cell suspensions were then collected and centrifuged at 300 x g for 5 min. at 4°C. Red blood cells were lysed using a lysing buffer (Life Technologies, Grand Island, NY). Next, the cells were washed with ice-cold 1X PBS and resuspended in 1X PBS for cell counting using an automatic cell counter (Cellometer Auto 2000; Nexcelom Bioscience, Lawrence, MA). Lastly, dead cells were removed from the cell suspensions using the Dead Cell Removal Kit (Miltenyi Biotec), and cells were counted again to determine final viable cell numbers.

#### Single-cell library preparation using the 10x Genomics platform

Viable cells were utilized for single-cell RNAseq library construction using the Chromium™ Controller and Chromium™ Single Cell 3’ Version 3 Kit (10x Genomics, Pleasanton, CA), following the manufacturer’s instructions. Briefly, viable cell suspensions were loaded into the Chromium™ Controller to generate gel beads in emulsion (GEM), with each GEM containing a single cell as well as barcoded oligonucleotides. Next, the GEMs were placed in the Veriti 96-well Thermal Cycler (Thermo Fisher Scientific, Wilmington, DE) and reverse transcription was performed in each GEM (GEM-RT). After the reaction, the complementary (c)DNA was cleaned using Silane DynaBeads (Thermo Fisher Scientific) and the SPRIselect Reagent Kit (Beckman Coulter, Indianapolis, IN). Next, the cDNA was amplified using the Veriti 96-well Thermal Cycler and cleaned using the SPRIselect Reagent Kit. Indexed sequencing libraries were then constructed using the Chromium™ Single Cell 3’ Version 3 Kit, following the manufacturer’s instructions.

cDNA was fragmented, end-repaired, and A-tailed using the Chromium™ Single Cell 3’ Version 3 Kit, following the manufacturer’s instructions. Next, adaptor ligation was performed using the Chromium™ Single Cell 3’ Version 3 Kit, followed by post-ligation clean-up using the SPRIselect Reagent Kit to obtain the final library constructs, which were then amplified using PCR. After performing a post-sample index double-sided size selection using the SPRIselect Reagent Kit, the quality and quantity of the DNA were analyzed using the Agilent Bioanalyzer High Sensitivity Chip (Agilent Technologies, Wilmington, DE). The Kapa DNA Quantification Kit for Illumina® platforms (Kapa Biosystems, Wilmington, MA) was used to quantify the DNA libraries, following the manufacturer’s instructions.

#### Single-nuclei sample preparation

We developed a new protocol to isolate nuclei from frozen placenta samples, based on DroNc-seq [94] and an early version of the protocol developed by the Martelotto lab [95]. For each placenta sample, 1mm frozen placenta biopsy punches were collected and immediately lysed with ice-cold lysis buffer (10 mM Tris-HCl, pH 7.5, 10 mM NaCl, 3 mM MgCl2, 2% BSA, 0.2 U/µl Protector RNase Inhibitor, and 0.1% IGEPAL-630) for 5 minutes. During incubation the samples were gently mixed by swirling the tube twice and collected by centrifugation at 500 x g for 5 minutes at 4°C. The process was repeated twice for a total of 3 cycles of lysis (5 minutes long each). Next, the pellets were washed with ice-cold nuclei suspension buffer (1X PBS containing 2% BSA and 0.2 U/µl Protector RNase Inhibitor, ROCHE) and filtered through a 30μm cell strainer (Fisher Scientific). Nuclei suspensions were then collected and centrifuged at 500 x g for 5 minutes at 4°C. Nuclei were counted using a Countess™ II FL (Thermo Fisher Scientific, Durham, NC). All samples exhibited 100% cell death with DAPI staining, indicative of complete cell lysis. Nuclei were then utilized for single-nuclei RNAseq library construction using the Chromium™ Controller and Chromium™ Single Cell 3’ version 2 kit (10x Genomics), following the manufacturer’s instructions.

#### Single-nuclei 10x Genomics library preparation

Briefly, nuclei suspensions for pools of 16 samples were loaded on the Chromium™ Controller to generate GEMs, with each GEM containing a single cell as well as barcoded oligonucleotides. Each pool was loaded on two different channels, for a total of two pools with two replicates each (4 libraries, 32 pregnancy cases). Next, the GEMs were placed in the Veriti 96-well Thermal Cycler, and reverse transcription was performed in each GEM (GEM-RT). After the reaction, the complementary (c)DNA was cleaned using Silane DynaBeads and the SPRIselect Reagent Kit. Next, the cDNA was amplified using the Veriti 96-well Thermal Cycler and cleaned using the SPRIselect Reagent kit. cDNA quality and quantity were analyzed using the Agilent Bioanalyzer High Sensitivity chip. cDNA was fragmented, end-repaired, and A-tailed using the Chromium™ Single Cell 3’ version 2 kit, following the manufacturer’s instructions. Next, adapter ligation was performed using the Chromium™ Single Cell 3’ version 2 kit followed by post-ligation cleanup using the SPRIselect Reagent kit to obtain the final library constructs, which were then amplified using PCR. After performing a post-sample index double-sided size selection using the SPRIselect Reagent kit, the quality and quantity of the DNA were analyzed using the Agilent Bioanalyzer High Sensitivity chip. The Kapa DNA Quantification Kit for Illumina® platforms was used to quantify the DNA libraries. Phix (Illumina, San Diego, CA), serially diluted at different concentrations, was used as the standard together with a negative and a positive control to quantify and normalize the libraries for loading of the sequencer.

#### Sequencing

Libraries were sequenced on the Illumina NextSeq 500 in the Luca/Pique-Regi laboratory and in the CMMG Genomics Services Center (GSC). The Illumina 75 Cycle Sequencing Kit was used with 58 cycles for R2, 26 for R1, and 8 for I1.

### scRNA-seq and snRNA-seq data analyses

Raw fastq files were downloaded from previously established resources (as detailed in “Data Availability”), and the new sequencing data were processed using Cell Ranger version 3.0.0 from 10X Genomics for de-multiplexing. The fastq files were then aligned using kallisto [96], and bustools [97] summarized the cell/gene transcript counts in a matrix for each sample, using the “lamanno” workflow for scRNA-seq and the “nucleus” workflow for snRNA-seq. Each sample was then processed using DIEM [98] to eliminate debris and empty droplets for both scRNA-seq and snRNA-seq. To avoid the loss of cells that may express viral receptors, we did not exclude cell doublets from the analyses included in this report, which should have negligible effects on the results and conclusions. All count data matrices were then normalized and combined using the “NormalizeData,” “FindVariableFeatures,” and “ScaleData” methods implemented in the Seurat package in R (Seurat version 3.1, R version 3.6.1) [99] and [100]. Afterward, the Seurat “RunPCA” function was applied to obtain the first 50 principal components, and the different batches and locations were integrated and harmonized using the Harmony package in R [101]. The top 30 harmony components were then processed using the Seurat “runUMAP” function to embed and visualize the cells in a two-dimensional map via the Uniform Manifold Approximation and Projection for Dimension Reduction (UMAP) algorithm [102, 103]. To label the cells, the Seurat “FindTransferAnchors” and “TransferData” functions were used for each group of locations separately to assign a cell-type identity based on our previously labeled data as reference panel (as performed in [61]). Cell type abbreviations used are: STB, Syncytiotrophoblast; EVT, Extravillous trophoblast; CTB, cytotrophoblast; HSC, hematopoietic stem cell; npiCTB, non proliferative interstitial cytotrophoblast; LED, lymphoid endothelial decidual cell. Visualization of viral receptor gene expression was performed using the ggplot2 [104] package in R with gene expression values scaled to transcripts per million (TPM).

### Bulk Gene Expression Data Analysis of ACE2 and TMPRSS2 in the placental tissues

Gene expression data for the study by Kim et al. [67] was available from the www.ebi.ac.uk/microarray-as/ae/ database (entry ID: E-TABM-577), while data for the study by Toft et al. [68] is available in our data repertoire. The mas5calls function from the *affy* package in Bioconductor was used to determine presence above background of each probeset corresponding to a given gene [105].

## Acknowledgments

We thank the physicians, nurses, and research assistants from the Center for Advanced Obstetrical Care and Research, the Intrapartum Unit, and the PRB Clinical Laboratory for their help with collecting and processing samples.

## Supplementary Material

**Figure S1.**
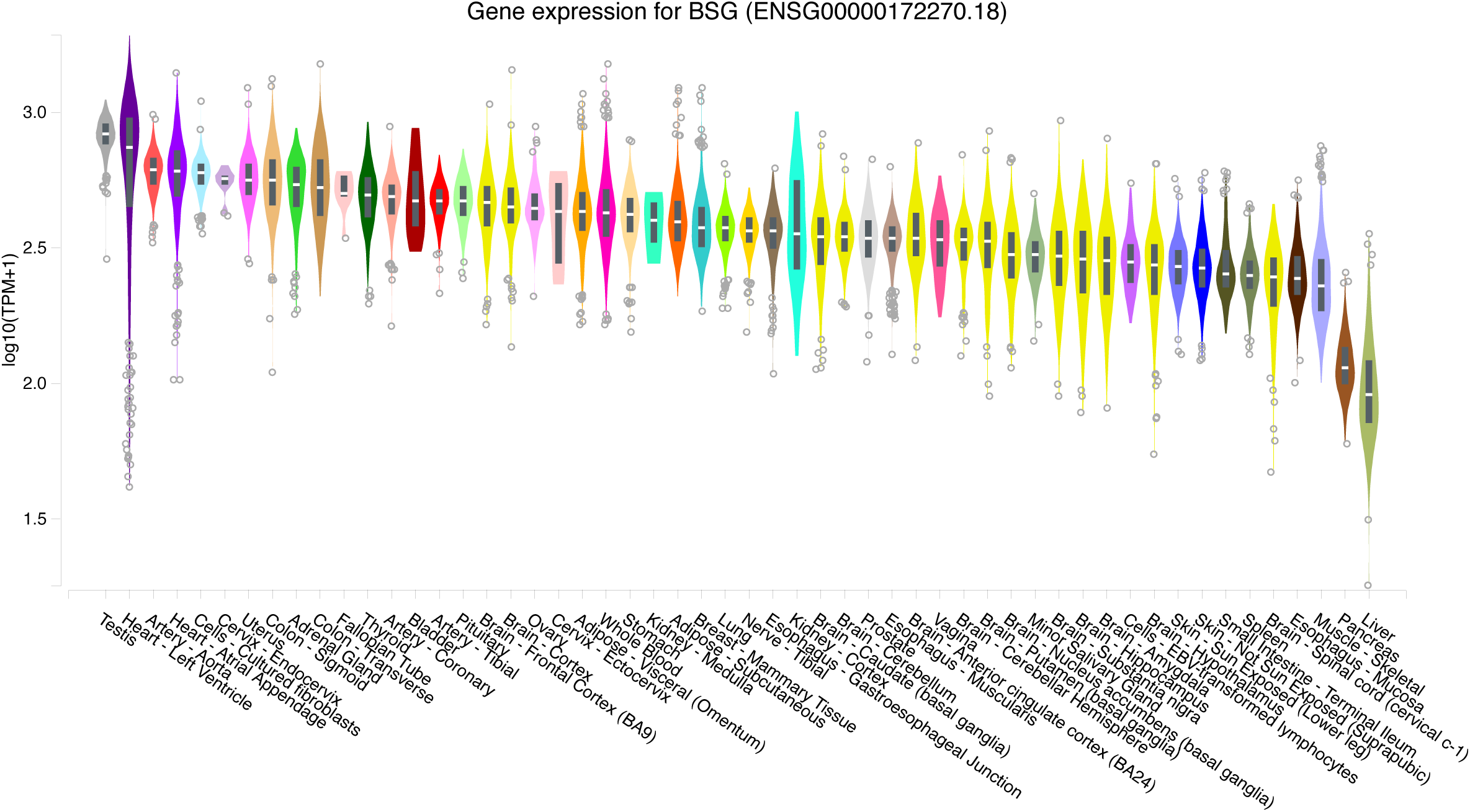
Gene expression values for BSG across tissues collected by the GTEx project. For each tissue, a violin plot and a boxplot describe the variation in gene expression levels for BSG across all of the samples collected by the project sorted left to right by median gene expression (plot generated through the GTEx portal).

**Figure S2.**
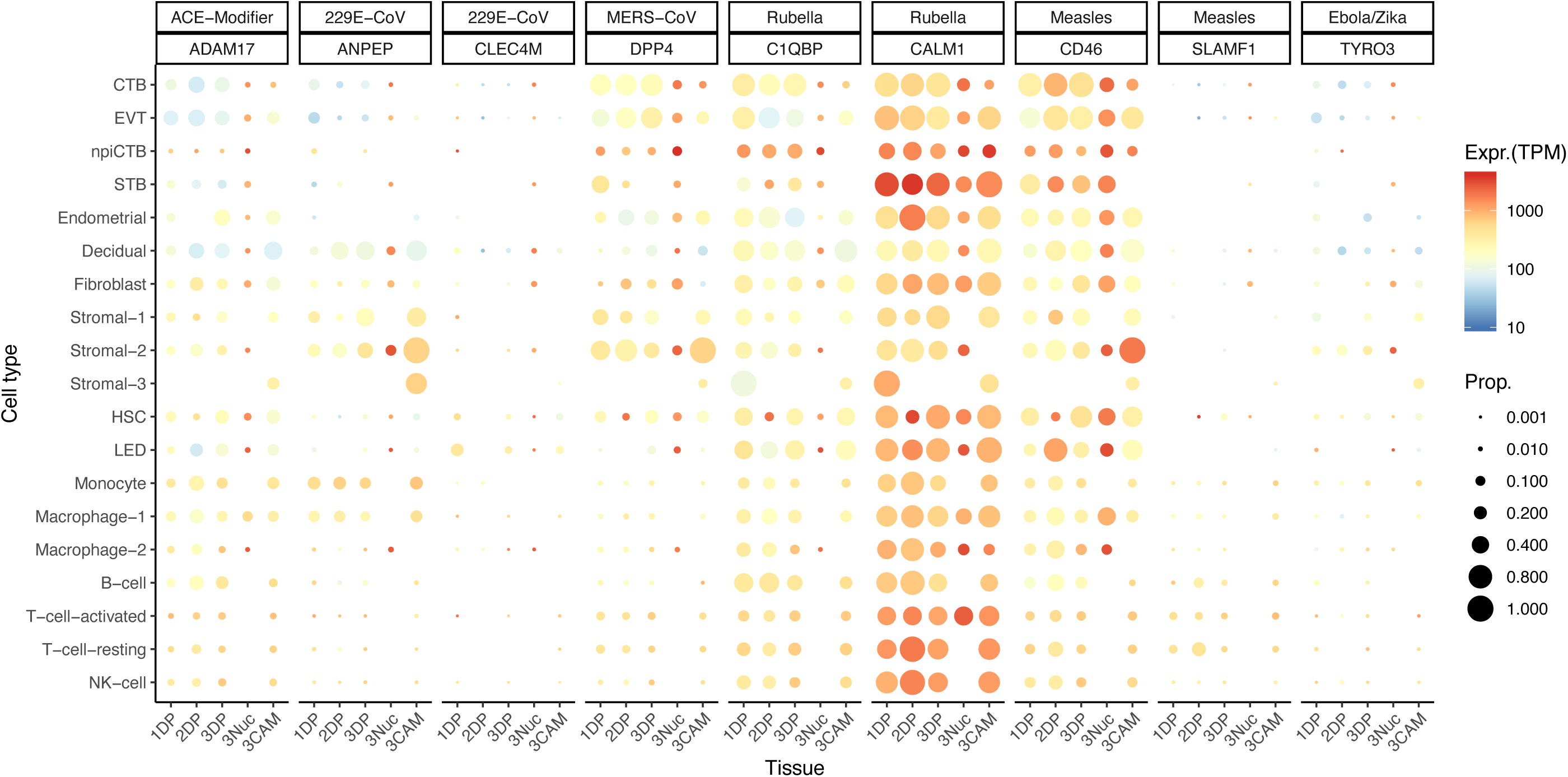
Dot plot depicting the expression of different viral receptors/molecules used by virus that caused congenital infection. Each row represents a different cell type, and columns are grouped first by virus type, receptor/molecule gene, and placental tissue/time-of sampling (1DP, 2DP and 3DP represent the first, second and third trimester, 3Nuc represents the third trimester nuclei, and 3CAM represents the third trimester chorioamniotic membranes). The size of the dot represents the proportion of cells that express the receptor with more than zero transcripts, and the color represents the average gene expression for the subset of cells expressing that gene in transcripts per million (TPM). Cell type abbreviations used are: STB, Syncytiotrophoblast; EVT, Extravillous trophoblast; CTB, cytotrophoblast; HSC, hematopoietic stem cell; npiCTB, non proliferative interstitial cytotrophoblast; LED, lymphoid endothelial decidual cell.

**Table S1:** Summary of all the single cell resources analyzed using existing and new data.

| Library ID | Type | Study | Location | Location description | #Pooled | #Cells | Avg. #UMI | Avg. #Genes |
| --- | --- | --- | --- | --- | --- | --- | --- | --- |
| HPL20289_R_PVBP_1 | scRNA-seq 10X 3' v3 | New, this study | 2DP | 2nd trim. Decidua + Placenta | 1 | 6340 | 9900 | 2057 |
| HPL20289_R_PVBP_2 | scRNA-seq 10X 3' v3 | New, this study | 2DP | 2nd trim. Decidua + Placenta | 1 | 6850 | 9810 | 1910 |
| CC7-1 | snRNA-seq 10X 3' v2 | New, this study | 3Nuc | 3rd trim. Dec. + Placenta Nuclei | 16 | 2454 | 1111 | 473 |
| CC7-2 | snRNA-seq 10X 3' v2 | New, this study | 3Nuc | 3rd trim. Dec. + Placenta Nuclei | 16 | 2759 | 1199 | 519 |
| CC8-1 | snRNA-seq 10X 3' v2 | New, this study | 3Nuc | 3rd trim. Dec. + Placenta Nuclei | 16 | 3831 | 1102 | 530 |
| CC8-2 | snRNA-seq 10X 3' v2 | New, this study | 3Nuc | 3rd trim. Dec. + Placenta Nuclei | 16 | 4267 | 1098 | 518 |
| s1DB | scRNA-seq 10X 3' v2 | Pique-Regi et al. (2019) | 3DP | 3rd trim. Decidua + Placenta | 1 | 2537 | 6560 | 1381 |
| s2DB | scRNA-seq 10X 3' v2 | Pique-Regi et al. (2019) | 3DP | 3rd trim. Decidua + Placenta | 1 | 2285 | 3695 | 967 |
| s2P | scRNA-seq 10X 3' v2 | Pique-Regi et al. (2019) | 3DP | 3rd trim. Decidua + Placenta | 1 | 2448 | 5475 | 1431 |
| s3DB | scRNA-seq 10X 3' v2 | Pique-Regi et al. (2019) | 3DP | 3rd trim. Decidua + Placenta | 1 | 2321 | 4477 | 1061 |
| s3P | scRNA-seq 10X 3' v2 | Pique-Regi et al. (2019) | 3DP | 3rd trim. Decidua + Placenta | 1 | 2412 | 9160 | 2047 |
| s4DB | scRNA-seq 10X 3' v2 | Pique-Regi et al. (2019) | 3DP | 3rd trim. Decidua + Placenta | 1 | 3064 | 6124 | 1539 |
| s4P | scRNA-seq 10X 3' v2 | Pique-Regi et al. (2019) | 3DP | 3rd trim. Decidua + Placenta | 1 | 2877 | 8196 | 2003 |
| s5DB | scRNA-seq 10X 3' v2 | Pique-Regi et al. (2019) | 3DP | 3rd trim. Decidua + Placenta | 1 | 4486 | 3644 | 950 |
| s5P | scRNA-seq 10X 3' v2 | Pique-Regi et al. (2019) | 3DP | 3rd trim. Decidua + Placenta | 1 | 2961 | 4847 | 1280 |
| s6DB | scRNA-seq 10X 3' v2 | Pique-Regi et al. (2019) | 3DP | 3rd trim. Decidua + Placenta | 1 | 3618 | 7034 | 1600 |
| s6P | scRNA-seq 10X 3' v2 | Pique-Regi et al. (2019) | 3DP | 3rd trim. Decidua + Placenta | 1 | 1228 | 10924 | 2359 |
| s7DB | scRNA-seq 10X 3' v2 | Pique-Regi et al. (2019) | 3DP | 3rd trim. Decidua + Placenta | 1 | 2737 | 8983 | 1811 |
| s7P | scRNA-seq 10X 3' v2 | Pique-Regi et al. (2019) | 3DP | 3rd trim. Decidua + Placenta | 1 | 2479 | 11049 | 2159 |
| s8DB | scRNA-seq 10X 3' v2 | Pique-Regi et al. (2019) | 3DP | 3rd trim. Decidua + Placenta | 1 | 2986 | 6265 | 1404 |
| s8P | scRNA-seq 10X 3' v2 | Pique-Regi et al. (2019) | 3DP | 3rd trim. Decidua + Placenta | 1 | 3344 | 12233 | 2588 |
| s9DB | scRNA-seq 10X 3' v2 | Pique-Regi et al. (2019) | 3DP | 3rd trim. Decidua + Placenta | 1 | 3916 | 5615 | 1243 |
| s1W | scRNA-seq 10X 3' v2 | Pique-Regi et al. (2019) | 3CAM | 3rd trim. Chorionamnionic membranes | 1 | 2921 | 5342 | 1467 |
| s2W | scRNA-seq 10X 3' v2 | Pique-Regi et al. (2019) | 3CAM | 3rd trim. Chorionamnionic membranes | 1 | 2836 | 6397 | 1750 |
| s3W | scRNA-seq 10X 3' v2 | Pique-Regi et al. (2019) | 3CAM | 3rd trim. Chorionamnionic membranes | 1 | 1961 | 9591 | 1914 |
| s4W | scRNA-seq 10X 3' v2 | Pique-Regi et al. (2019) | 3CAM | 3rd trim. Chorionamnionic membranes | 1 | 2586 | 6672 | 1645 |
| s5W | scRNA-seq 10X 3' v2 | Pique-Regi et al. (2019) | 3CAM | 3rd trim. Chorionamnionic membranes | 1 | 1966 | 4668 | 1266 |
| s6W | scRNA-seq 10X 3' v2 | Pique-Regi et al. (2019) | 3CAM | 3rd trim. Chorionamnionic membranes | 1 | 4813 | 1682 | 592 |
| s7W | scRNA-seq 10X 3' v2 | Pique-Regi et al. (2019) | 3CAM | 3rd trim. Chorionamnionic membranes | 1 | 3207 | 4507 | 1175 |
| s8W | scRNA-seq 10X 3' v2 | Pique-Regi et al. (2019) | 3CAM | 3rd trim. Chorionamnionic membranes | 1 | 5037 | 7069 | 1708 |
| s9W | scRNA-seq 10X 3' v2 | Pique-Regi et al. (2019) | 3CAM | 3rd trim. Chorionamnionic membranes | 1 | 3938 | 7483 | 1589 |
| FCA7167219 | scRNA-seq 10X 3' v2 | Vento-Tormo et al. (2018) | 1DP | 1st trim. Decidua + Placenta | 1 | 716 | 3591 | 836 |
| FCA7167221 | scRNA-seq 10X 3' v2 | Vento-Tormo et al. (2018) | 1DP | 1st trim. Decidua + Placenta | 1 | 1252 | 3607 | 941 |
| FCA7167222 | scRNA-seq 10X 3' v2 | Vento-Tormo et al. (2018) | 1DP | 1st trim. Decidua + Placenta | 1 | 1928 | 9790 | 1872 |
| FCA7167223 | scRNA-seq 10X 3' v2 | Vento-Tormo et al. (2018) | 1DP | 1st trim. Decidua + Placenta | 1 | 8080 | 2732 | 895 |
| FCA7167224 | scRNA-seq 10X 3' v2 | Vento-Tormo et al. (2018) | 1DP | 1st trim. Decidua + Placenta | 1 | 2232 | 6132 | 1675 |
| FCA7167226 | scRNA-seq 10X 3' v2 | Vento-Tormo et al. (2018) | 1DP | 1st trim. Decidua + Placenta | 1 | 5775 | 7415 | 1918 |
| FCA7196218 | scRNA-seq 10X 3' v2 | Vento-Tormo et al. (2018) | 1DP | 1st trim. Decidua + Placenta | 1 | 9028 | 2203 | 828 |
| FCA7196219 | scRNA-seq 10X 3' v2 | Vento-Tormo et al. (2018) | 1DP | 1st trim. Decidua + Placenta | 1 | 3704 | 8209 | 1971 |
| FCA7196220 | scRNA-seq 10X 3' v2 | Vento-Tormo et al. (2018) | 1DP | 1st trim. Decidua + Placenta | 1 | 7338 | 6139 | 1862 |
| FCA7196224 | scRNA-seq 10X 3' v2 | Vento-Tormo et al. (2018) | 1DP | 1st trim. Decidua + Placenta | 1 | 7930 | 4028 | 1182 |
| FCA7196225 | scRNA-seq 10X 3' v2 | Vento-Tormo et al. (2018) | 1DP | 1st trim. Decidua + Placenta | 1 | 4121 | 6144 | 1511 |
| FCA7196226 | scRNA-seq 10X 3' v2 | Vento-Tormo et al. (2018) | 1DP | 1st trim. Decidua + Placenta | 1 | 3380 | 15098 | 2564 |
| FCA7474062 | scRNA-seq 10X 3' v2 | Vento-Tormo et al. (2018) | 1DP | 1st trim. Decidua + Placenta | 1 | 2149 | 6351 | 1523 |
| FCA7474063 | scRNA-seq 10X 3' v2 | Vento-Tormo et al. (2018) | 1DP | 1st trim. Decidua + Placenta | 1 | 2715 | 15947 | 2703 |
| FCA7474064 | scRNA-seq 10X 3' v2 | Vento-Tormo et al. (2018) | 1DP | 1st trim. Decidua + Placenta | 1 | 4941 | 13354 | 2889 |
| FCA7474065 | scRNA-seq 10X 3' v2 | Vento-Tormo et al. (2018) | 1DP | 1st trim. Decidua + Placenta | 1 | 4878 | 13479 | 2855 |
| FCA7474068 | scRNA-seq 10X 3' v2 | Vento-Tormo et al. (2018) | 1DP | 1st trim. Decidua + Placenta | 1 | 4298 | 15640 | 2534 |
| FCA7511881 | scRNA-seq 10X 3' v2 | Vento-Tormo et al. (2018) | 1DP | 1st trim. Decidua + Placenta | 1 | 2072 | 2214 | 530 |
| FCA7511882 | scRNA-seq 10X 3' v2 | Vento-Tormo et al. (2018) | 1DP | 1st trim. Decidua + Placenta | 1 | 2368 | 10295 | 1915 |
| FCA7511884 | scRNA-seq 10X 3' v2 | Vento-Tormo et al. (2018) | 1DP | 1st trim. Decidua + Placenta | 1 | 4534 | 9945 | 2090 |

**Table S2.** Clinical and demographic characteristics of the study population from which placental samples were collected for snRNAseq studies.

|  | **Study group**<br>**(n=32)** |
| --- | --- |
| **Clinical parameters** |  |
| Maternal age (years; median [IQR]) | 25 (21.8-29) |
| Body mass index (kg/m<sup>2</sup>; median [IQR]) | 27.8 (21.7-31.6)<sup>a</sup> |
| Primiparity | 15.6% (5/32) |
| Cesarean section | 59.4% (19/32) |
| Gestational age at delivery (weeks; median [IQR]) | 36.9 (32.9-39.1) |
| Birthweight (g; median [IQR]) | 2802.5 (1833.8-3233.8) |
| **Ethnicity** |  |
| African-American | 81.3% (26/32) |
| Caucasian | 6.2% (2/32) |
| Other | 12.5% (4/32) |

Data are given as median (interquartile range, IQR) and percentage (n/N)
<sup>a</sup> One missed data

**Table S3.** Bulk Gene Expression Data Analysis of ACE2 and TMPRSS2 in the placental tissues.

| **Study** | **Affymetrix probeset** | **Symbol** | **Log<sub>2</sub> Average Expression** | **% of samples detected above background** |
| --- | --- | --- | --- | --- |
| Kim et al. | 219962_at | ACE2 | 4.61 | 90% |
| Kim et al. | 222257_s_at | ACE2 | 5.71 | 70% |
| Kim et al. | 205102_at | TMPRSS2 | 5.97 | 10% |
| Kim et al. | 1570433_at | TMPRSS2 | 3.37 | 0% |
| Kim et al. | 211689_s_at | TMPRSS2 | 2.96 | 0% |
| Kim et al. | 226553_at | TMPRSS2 | 6.02 | 0% |
| Toft et al. | 219962_at | ACE2 | 4.26 | 82% |
| Toft et al. | 222257_s_at | ACE2 | 5.18 | 57% |
| Toft et al. | 1570433_at | TMPRSS2 | 3.30 | 0% |
| Toft et al. | 205102_at | TMPRSS2 | 5.78 | 0% |
| Toft et al. | 211689_s_at | TMPRSS2 | 3.34 | 0% |
| Toft et al. | 226553_at | TMPRSS2 | 5.65 | 0% |

